# Targeted systematic evolution of an RNA platform neutralizing DNMT1 function and controlling DNA methylation

**DOI:** 10.1101/2020.07.29.226803

**Authors:** CL Esposito, I Autiero, MA Basal, A Sandomenico, S Ummarino, M Borchiellini, M Ruvo, S Catuogno, AK Ebralidze, V de Franciscis, A Di Ruscio

## Abstract

DNA methylation is a fundamental epigenetic modification regulating gene expression. Aberrant DNA methylation is the most common molecular lesion in cancer cells. However, medical intervention has been limited to the use of toxic, unspecific demethylating drugs. Aptamers are novel high affinity targeting ligand molecules. By conjugating the inherent DNMT1 inhibiting capabilities of RNA to an aptamer platform, we generated a first-of-its kind aptamer approach that can target and neutralize DNMT1 function – the aptaDiR. Molecular modelling of RNA-DNMT1 complexes coupled with biochemical and cellular assays enabled the identification and characterization of aptaDiR. This novel RNA bio-drug blocks DNA methylation and impairs cancer cell viability.

Collectively, we present an innovative RNA-based approach to modulate DNMT1 activity in cancer or diseases characterized by aberrant DNA methylation and suggest the first alternative strategy to overcome the limitations of currently approved hypomethylating protocols, which will greatly improve clinical intervention on DNA methylation.

## Introduction

DNA methylation is a key epigenetic signature implicated in the regulation of gene expression ^1,2^. Methylation of CpG-rich (mCpG) promoters is carried out by members of the DNA methyltransferase (DNMT) family (DNMT1, DNMT2, DNMT3A, DNMT3B and DNMT3L) ^3,4^. Several studies have established a link between aberrant promoter methylation and cancer. Reduced DNA methylation contributes to genomic instability while site-specific aberrant methylation of gene promoters, i.e. tumour suppressor genes and transcription factors, results in specific gene silencing ^5,6^. Aberrant epigenetic modifications likely occur in the early stages of the tumour development and are reversible thereby offering the possibility to restore normal gene function abrogated by malignant cellular transformation. In light of this, “epigenetic targeting” brought about by specific and effective inhibitors could lead to clinically relevant strategies for cancer therapy ^7^.

Thus far, two nucleoside-based compounds, 5-azacytidine and 5-aza 2’-deoxy-cytidine, have been approved as DNMT inhibitors to reduce global DNA methylation levels. However, the lack of selectivity, toxicity and chemical instability has raised serious concerns for the use of these nucleoside analogues ^8-10^, revealing the clinical need to develop smarter and safer epigenetic drugs.

Nucleic-acid aptamers are a promising class of three-dimensional structured oligonucleotides that serve as high affinity ligands and potential antagonists of disease-associated proteins ^11^. They are usually selected from a large random sequence library through a combinatorial chemistry strategy named SELEX (Systematic Evolution of Ligands by EXponential enrichment) ^12^. Besides being cost-effective and relatively easy to manipulate, the small size of aptamers (6-30kD) enable them to access binding pockets that are usually inaccessible to macromolecules. In addition, they are not immunogenic and exhibit low toxicity, while retaining high affinity for the binding targets. These features make aptamers a valuable tool in clinical diagnosis, therapy and targeted delivery ^13,14^.

We previously discovered a class of RNAs able to inhibit DNMT1 enzymatic activity and regulate DNA methylation^15^. Specifically, we showed that a non-coding RNA, named *ec-CCAAT-enhancer-binding proteins alpha* (*CEBPA*), originating within *CEBPA* locus could interact with and inhibit DNMT1, thereby preventing methylation of *CEBPA* locus. Globally, we proved that the DNMT1 site-specific sequestration by DNMT1-interacting RNAs (DiRs) occurs at multiple loci and is dependent upon the presence of RNA stem-loop-like-secondary structures ^15^. Further, we identified the minimal 22 nucleotide-long recognition motif required to block DNMT1 activity ^15^, delineating an RNA-mediated mechanism controlling DNA methylation establishment. Herein, we adopt a doped SELEX approach to generate DNMT1-neutralizing RNA aptamers with enhanced affinity, specificity and stability than the natural existing counterparts. Through a combination of *in silico* modelling and *in vitro* biochemical and cellular-based assays, we demonstrate the applicability of this targeted DNMT1-aptamer platform, the aptaDiR, as the first RNA-based approach to correct aberrant DNA methylation and as a promising option for the treatment of cancer or other diseases characterized by aberrant DNA methylation.

## Results

### Evolution of anti-DNMT1 RNA aptamers

We have previously shown that a 22-nucleotide (nt)-long stem-loop-like RNA structure (R5), embedded in the sequence of the DNMT1-interacting RNA originating within the *CEBPA* locus, is sufficient to bind to the DNMT1 catalytic domain and inhibit its activity^15^. Specific tridimensional structures are involved in the formation of the RNA-DNMT1 complex ^15^, hence, we sought to model *in silico* the interaction between R5 or its mutant R5 (mutR5, which is unable to fold in secondary structures), with the human (h)DNMT1 catalytic region. The coordinates of the murine DNMT1 solved structure (PDB: 4d4a) in complex with its DNA substrate was used as a template (see methods section for details). This analysis revealed a different arrangement of R5 with respect to the mutR5. Indeed, while the former adopts a stem loop-like architecture that perfectly accommodates within the hDNMT1 catalytic region, thereby mimicking and competing out its natural DNA substrate, the latter adopts a conformation that hampers an adequate fitting within the DNMT1 binding site (**Figure 1**).

**Figure 1.**
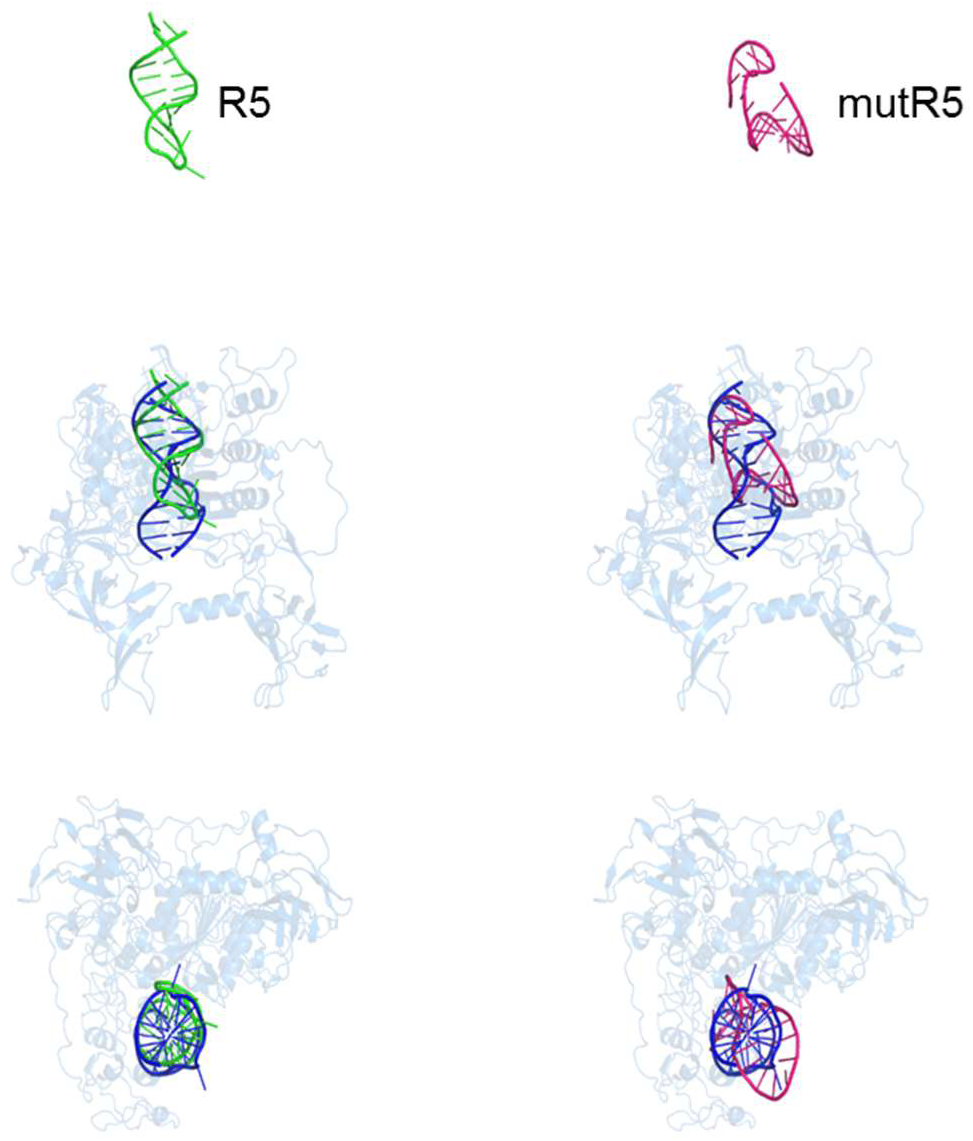
Models of R5 and mutR5. Cartoon representation of the tridimensional models of R5 (green) and mutR5 (magenta) and their complexes with the DNMT1 (blue) protein. The DNA substrate of the Xray structure used guide to build DNMT1 complexes is also shown in blue.

To evolve anti-DNMT1 RNA aptamers with increased affinity and stability, we built up a doped protein SELEX strategy by using as starting pool R5 variants with three fully randomized short regions (**Figure 2a** and **b** and Methods). The addition of 2’-Fluoro-Pyrimidines (2’F-Py) analogues at each position provided enhanced resistance to degradation of the RNA library. The 2’F-Py modified R5 (referred as DNMT1 bait) preserves its binding to DNMT1 as assessed by direct-Enzyme-Linked Oligonucleotides Assay (ELONA) (**Supplementary Figure 1**).

**Figure 2.**
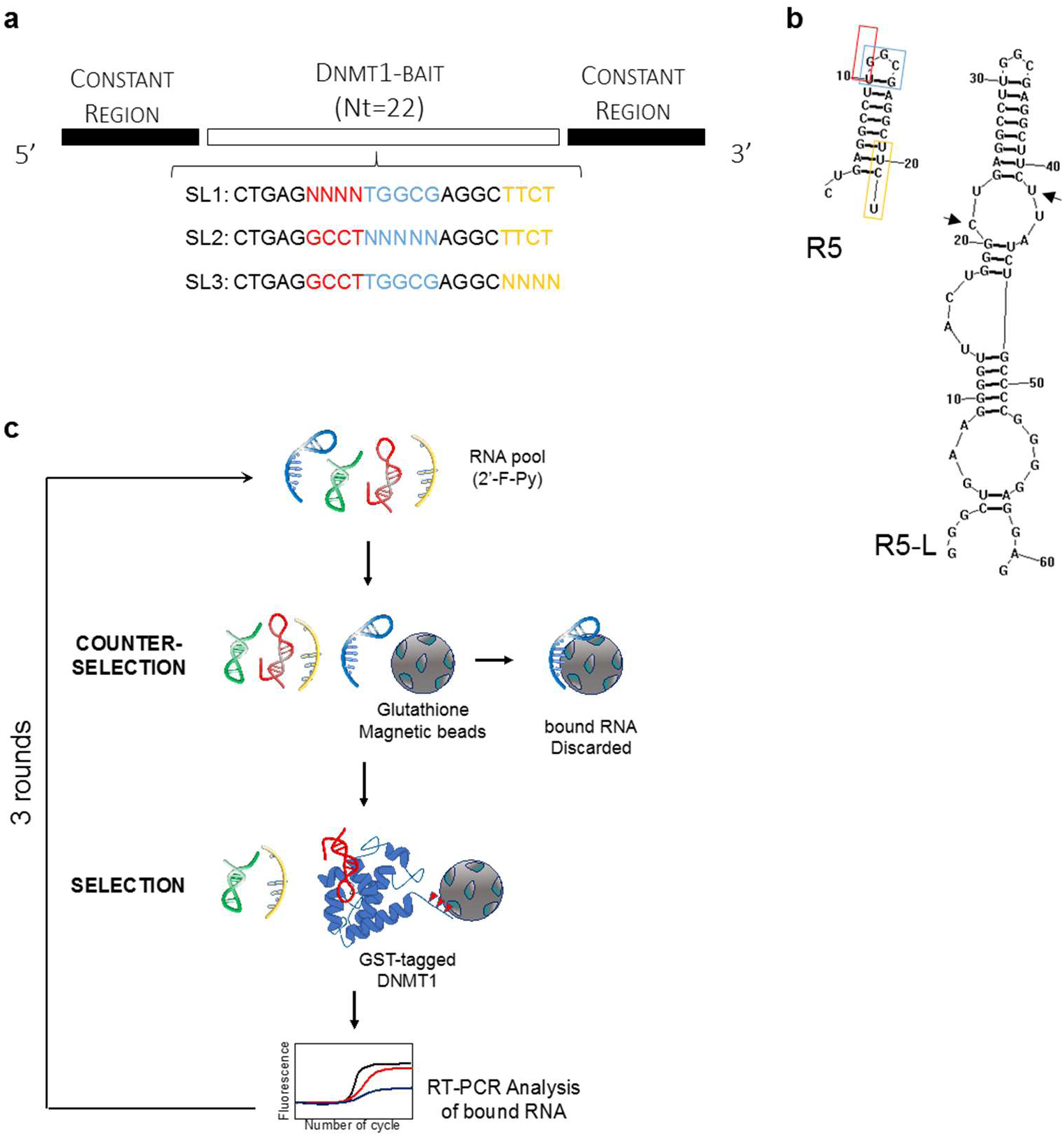
Selection process of anti-DNMT1 aptamers. (**a**) Diagram of the site-specific randomized sub-libraries (SL1, SL2 and SL3) used as starting pool for the SELEX cycles. (**b**) Stem– loop predicted structures of R5 and long R5 (R5-L). The black arrows indicate R5 sequence within R5-L. Regions within R5 that were randomized for the SELEX starting pool are boxed (SL1: red; SL2: light blue; SL3: yellow). (**c**) Scheme of the SELEX rounds. Each round include steps of: i) incubation of the RNA pool with glutathione magnetic beads (counter-selection); ii) recovering of unbound sequences; iii) incubation of the unbound sequences with purified GST-tagged DNMT1 protein (selection); iv) partitioning of the bound sequences with glutathione coupled magnetic beads; v) recovering and amplification of bound sequences by RT-PCR.

The SELEX library was designed starting from a construct (R5L) containing *ecCEBPA* R5 (DNMT1 bait) flanked by two constant 20- and 19-nt long naïve regions, at the 5’ and 3’ ends, respectively. The T7 promoter sequence engineered upstream of the 5’ end allowed transcription and amplification of the RNA construct (**Figure 2a**). Importantly, the R5L retained the stem-and-loop-like structure required for the DNMT1-RNA interaction (**Figure 2b**). Three different libraries (SL1, SL2 and SL3) were thus produced by degenerating three different regions (4-5 bases each) from the 5’, the central loop and the 3’ of the 22-nt DNMT1-bait (**Figure 2a** and **b**). The three libraries were mixed at an equimolar concentration and used as template for the transcription of the 2’F-Py modified RNA starting pool. The starting pool was then subjected to three rounds of SELEX using the purified Glutathione s-Transferase (GST)-tagged DNMT1 protein as the target. As schematized in **Figure 2c**, at each round, the pool was first incubated with glutathione coupled magnetic beads for a counter-selection step. The bound sequences were separated using a magnetic separator and the unbound aptamers were used for the selection on GST-tagged DNMT1. Following the selection step, bound sequences were partitioned on glutathione coupled magnetic beads and recovered by reverse transcription PCR (RT-PCR). During the three rounds, we used an increasing number of final washes to progressively enhance the stringency of the selection and to recover aptamers with higher affinity for the target (**Supplementary Table 1**).

### Selection of DNMT1-specific individual aptamers

After the third SELEX round, the final enriched pool was cloned using TA cloning system and approximately seventy clones isolated and sequenced. The sequences were aligned by *Muscle* algorithm and clustered into families. Among the analysed clones (not shown), variants coming from the three SLs were equally represented. The Ce-9 and Ce-10 aptamers were identified as the most enriched in the 3 clusters of sequences (**Supplementary Figure 2**), while a third aptamer Ce-49, was chosen from a separate cluster. DNMT1-aptamers binding was confirmed by ELONA. A binding efficiency comparable or even stronger than R5-L was detected for all analysed sequences in a range of 200 nM (**Figure 3a**). Efficient and cost-effective chemical synthesis is a key aspect of aptamer optimization. Thus, the initially identified 61-nt long aptamer sequences was reduced to an ideal 22/25-nt length, without altering the stem-loop-like structures critical for the interaction with DNMT1. The shortened (sh) aptamers: one for Ce-9 or Ce-49 (indicated as Ce-9 sh and Ce-49 sh, respectively) and two for Ce-10 (Ce10-1 sh, Ce10-2 sh), displayed the same hairpin-loop predicted structure as the longer sequences (**Figure 3b**) and preserved the binding ability to DNMT1 (**Figure 3c**), thus confirming the stem-loop-like structures to be sufficient for the aptamer-DNMT1 interaction.

**Figure 3.**
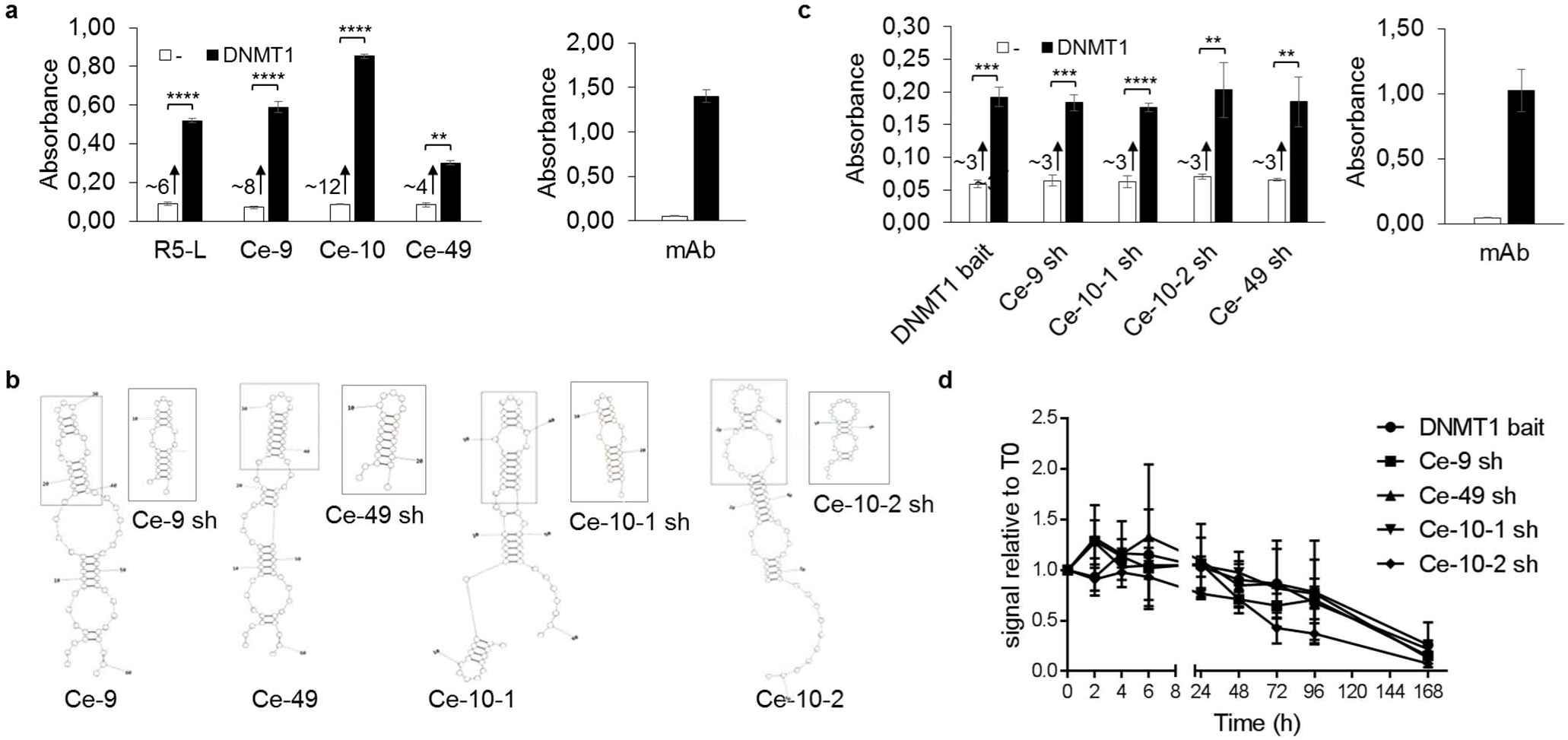
Analyses and optimization of individual aptamers from SELEX. (**a**) Binding of biotinylated selected aptamers from SELEX and R5-L on DNMT1 purified protein was analysed by ELONA (*left panel*). Anti-DNMT1 antibody (mAb, *right panel*) was used as a positive control. Error bars depict mean ± SD (n=3). Statistics by *t*-test: **, p<0.01; ****, p<0.0001. Fold increase of binding is reported. (**b**) Predicted secondary structures of Ce-49, Ce-9 and Ce-10 and designed shortened aptamers (boxed). Two different folding and correspondent short aptamers were predicted for Ce-10 and indicated as Ce-10-1 and Ce-10-2, respectively. The portions of the short versions within the corresponding long aptamer are boxed. (**c**) Binding of 3’-biotinylated short aptamers and DNMT1 bait on DNMT1 purified protein was analysed by ELONA. Error bars depict mean ± SD (n=3). Statistics by *t*-test: **, p<0.01; ***, p<0.001; ****, p<0.0001. Anti-DNMT1 antibody (mAb, *right*) was used as a positive control. Fold increase of binding is reported. (**d**) Short aptamers and DNMT1 bait serum stability were measured in 85% human serum for indicated times. At each time point, RNA-serum samples were collected and evaluated by electrophoresis with 15% denaturing polyacrylamide gel. The intensity of the bands was quantified and expressed relative to T0.

We then analysed the stability of DNMT1-bait vis-à-vis the selected aptamers when incubated with high concentrations of human serum (85%) at 37 °C, over an extended time. The four aptamers and the DNMT1-bait remain almost undigested up-to 48 hours, owing to the presence of the 2’F Py modification (**Figure 3d** and **Supplementary Figure 3**).

In summary, our approach led to the identification of four very stable short RNA aptamer ligands for DNMT1.

### Characterization of binding and target specificity of the DNMT1 aptamers

To probe the biomolecular interactions and test by direct binding, the aptamer affinity to DNMT1, we took advantage of the Bio-Layer Interferometry technology ^16^ implemented in the BLItz instrument. Among the tested sequences, the best dose-dependent binding to the immobilized DNMT1 protein was detected for Ce-49 sh and Ce-10-2 sh (**Figure 4a** and **b**). The affinity measurement (**Supplementary Figure 4a** and **b**) provided KD values of 79.2 ± 23.7*10^−9^ M (R2 =0.9646) and 66.4 ± 24.0 *10^−9^ M (R2 =0.9426) for Ce-49 sh and Ce-10-2 sh, respectively. Dose-response binding analyses were also performed with the DNMT1-bait at concentrations ranging between 100 nM and 2 μM, since no binding was detected at lower concentrations (**Figure 4c**). Data fitting (**Supplementary Figure 4c**) provided a KD value of 0.6 ± 0.1 *10^−6^ M (R2 =0.9017), about 10 times higher as compared to that of the new molecules. The binding curves also showed stronger signals (about 1.25 and 1.80 nm, respectively, at the highest concentration) for Ce-49 sh and Ce-10-2 sh with respect to DNMT1 bait (about 0.17 nm), a difference that further affirms the increased affinity of the two newly generated aptamers (**Figure 4a-c**). In accordance with previous findings ^15^, no binding was detected for the mutR5 sequence which is unable to fold in stem-loop-like structures and used as a negative control (**Figure 4d**). The improved binding capability of the selected aptamers was further validated by RNA Electrophoretic Mobility-Shift Assay (REMSA) (**Figure 4e**). A fundamental aspect for the use *in vivo* of aptamers selected with recombinant proteins is to show their ability to bind the native target protein in the proper cellular context. Therefore, to check aptamer binding to the endogenous DNMT1 protein we carried out an aptamer-mediated pull-down assay. Extracts from K562 leukemia cell model, showing high levels of DNMT1, were incubated with biotin-tagged DNMT1 bait, Ce-49 sh or Ce-10-2 sh and the complexes were purified on streptavidin-coated beads, followed by immunoblotting with anti-DNMT1 antibody. All tested sequences display interaction with the native DNMT1 (**Figure 4f**).

**Figure 4.**
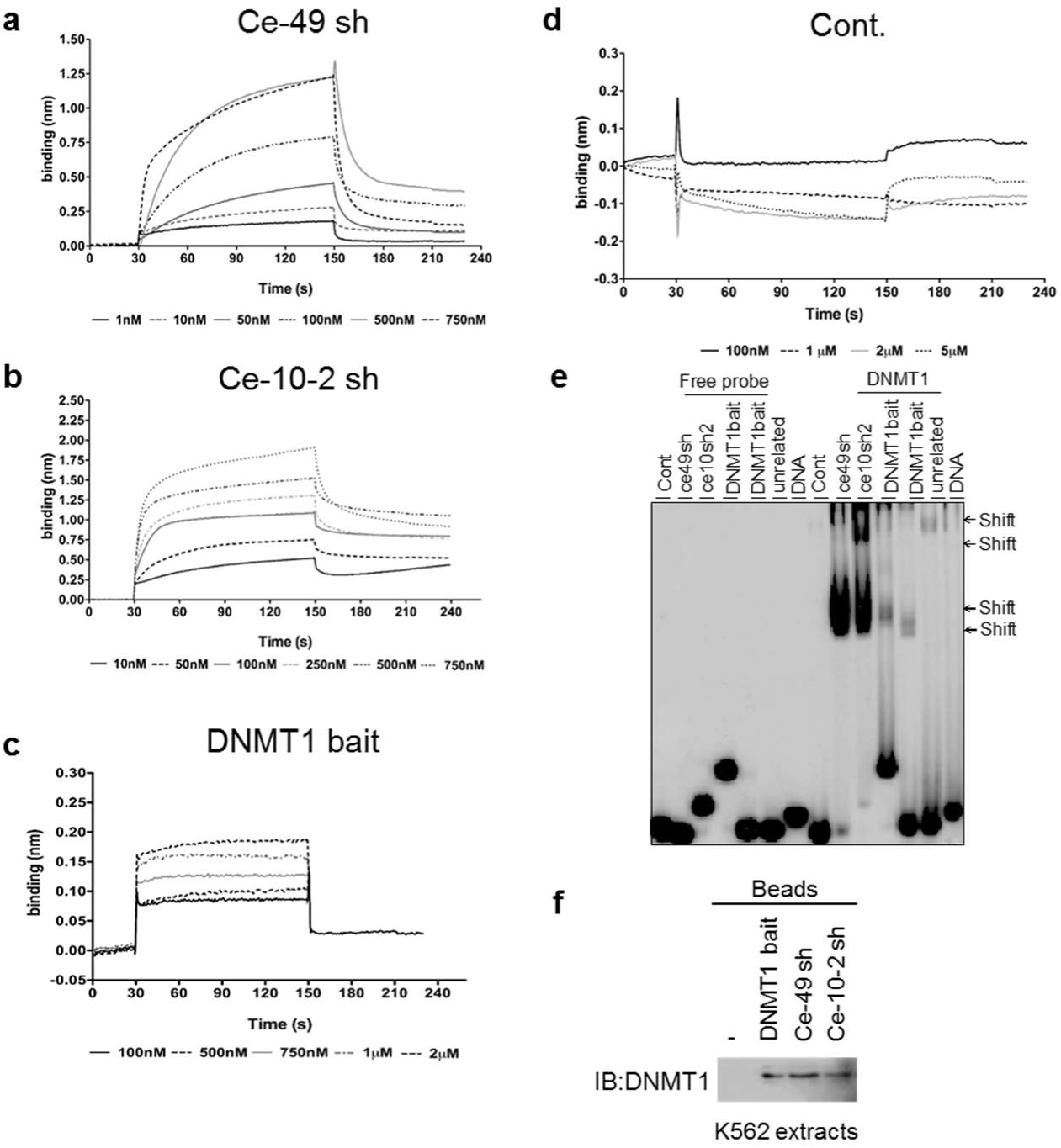
Aptamer affinity and *in vivo* binding to DNMT1. (**a**-**d**) Bio-Layer Interferometry dose-response measurements of Ce-49 sh (**a**), Ce-10-2 sh (**b**), DNMT1 bait (**c**) or mut-R5 (Cont., used as a negative control) binding to DNMT1 functionalized-biosensors (**d**). (**e**) The interaction of DNMT1 and DNMT1 bait or indicated aptamers was analysed by EMSA. (**f**) Aptamer-mediated pulldown of DNMT1. Protein extract from K562 cells were incubated with the biotinylated DNMT1 bait, Ce-49 sh and Ce-10-2 sh. Bound proteins were purified on streptavidin beads and immunoblotted with anti-DNMT1 antibodies.

As a following step we evaluated the DNMT1 selectivity and specificity of the selected aptamers Ce-49 sh and Ce-10-2 sh as compared to the DNMT1-bait, by performing comparative binding experiments with the other main members of the DNMT family: DNMT3A and DNMT3B and with the unrelated chromatin modifier lysine acetyltransferase 5 (KAT5). No significant interaction was recorded between the tested aptamers (**Figure 5a, b**), that show the same specificity as DNMT1 bait, and the three control proteins (**Figure 5c**). Finally, we tested the binding of the aptamers to the Human serum albumin (HSA) the most abundant plasma protein. HSA binds nucleic acids in a nonspecific manner through its positive charges and may reduce the bioavailability of circulating aptamers, thereby limiting their clinical applicability. Remarkably, no interaction was detected by incubating increasing concentrations up to 750 nM of Ce-49 sh and Ce-10-2 sh, with HSA (**Figure 5d** and **Supplementary Figure 5**).

**Figure 5.**
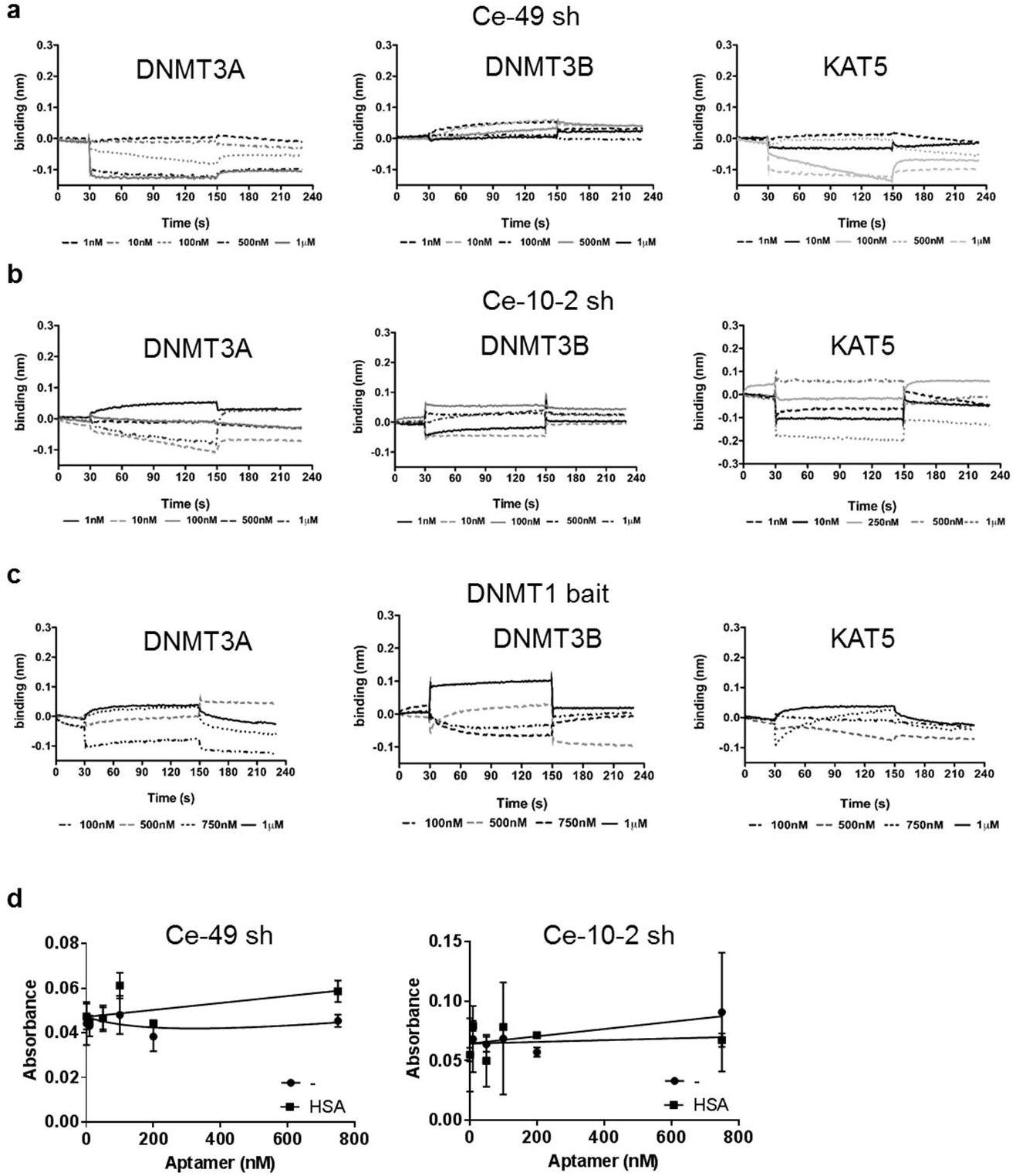
Aptamer specificity. (**a-c**) Binding measured by bio-layer interferometry of Ce-49 sh (**a**), Ce-10-2 sh (**b**) and DNMT1 bait (**c**) to DNMT3A (*left panels*), DNMT3B (*middle panels*) and KAT5 protein (*right panels*) immobilized on separate biosensors. Aptamers were tested at the reported concentrations. (**d**) ELONA assay of indicated biotinylated aptamers on plates left uncoated (-) or coated with HSA purified protein.

Taken together, these data show that Ce-49 sh and Ce-10-2 sh are high affinity and specific ligands for DNMT1 with apparent *in vitro* KDs within the nanomolar range, an appealing feature to proceed to clinical studies.

### Molecular Dynamic simulations of DNMT1-aptamer complexes

To dissect the structural and dynamic properties of DNMT1 interaction with the selected aptamers, we conducted *in silico* molecular simulations in explicit waters for 300 nanoseconds (ns) with the parental R5–, Ce-49 sh– or Ce-10-2 sh–DNMT1 complexes, respectively (**Figure 6**). The values of the root mean square deviation (RMSD) (**Supplementary Figure 6**) of the trajectory structures versus the starting models indicate stability of the system during the entire simulation with slightly dissimilar behaviour of the Ce-10-2 sh–DNMT1. In Ce10-2sh–DNMT1, the RMSD values exhibited by either the complex or each individual component were higher to some degree than those observed in R5-DNMT1 and Ce-49 sh–DNMT1 complexes (see **Supplementary Figure 6** and **7** for details). Additionally, the number of persistent hydrogen bonds within the last 150 ns simulation (**Supplementary Figure 8**) resulted more stable at the aptamer-protein interfaces for the R5 and Ce-49 sh complexes than the Ce-10-2 sh-DNMT1 interface. These findings suggest that Ce-10-2 sh– DNMT1 complex undergoes to greater rearrangement in the time scale used, revealing important structural determinants related to the aptamer sequence and responsible for the stability of the aptamer complexes with the target.

**Figure 6.**
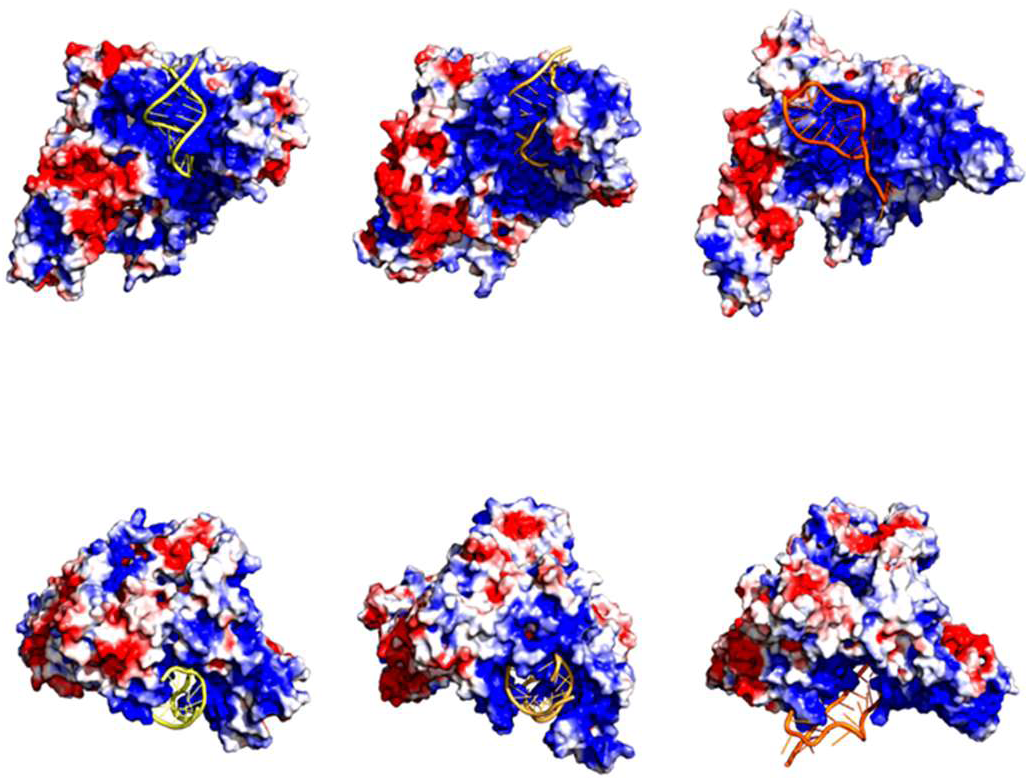
Molecular dynamic analyses. Molecular dynamic representative structures of R5 or Ce-49 and Ce-10-2 sh aptamer-DNMT1 complex simulations derived using a RMSD based clustering approach. DNMT1 protein is shown in surface and coloured according to the electrostatic potential. The red colour (negative potential) arises from an excess of negative charges near the surface and the blue colour (positive potential) occurs when the surface is positively charged (± 3 kT/e). The white regions correspond to fairly neutral potentials. For aptamer regions, the following colour scheme was adopted: R5: yellow, Ce-49 sh: brown and Ce-10-2 sh: orange.

Collectively, these results validate the complex stability and confirm a similar binding modality between the selected aptamers and the parental sequence with DNMT1.

### DNMT1-specific aptamer functional characterization

To examine the ability of the selected aptamers (Ce-49 sh and Ce-10-2 sh) to inhibit DNMT1-mediated methylation, we performed *in vitro* DNMT1 inhibition assay. The enzymatic activity of DNMT1 purified protein was measured in the absence or presence of DNMT1-bait or the selected aptamers. All the sequences blocked DNMT1 as shown by the 50% reduction of its activity (**Figure 7a**). Consistently, a reduction of 40 to 60% DNMT1 activity was reached when testing nuclear extracts obtained from K562 cells previously transfected with the Ce-49 sh and Ce-10-2 sh, as compared to the untreated cells (**Figure 7b**). As expected, no effect was observed using extracts from cells transfected with mutR5 (indicated as Cont.). Further, the DNMT1-bait and Ce-49 sh and Ce-10-2 sh were unable to interfere *in vitro* with the enzymatic activity of DNMT3A/B (**Figure 7c**), thus confirming the high affinity and selectivity of the aptamers for DNMT1.

**Figure 7.**
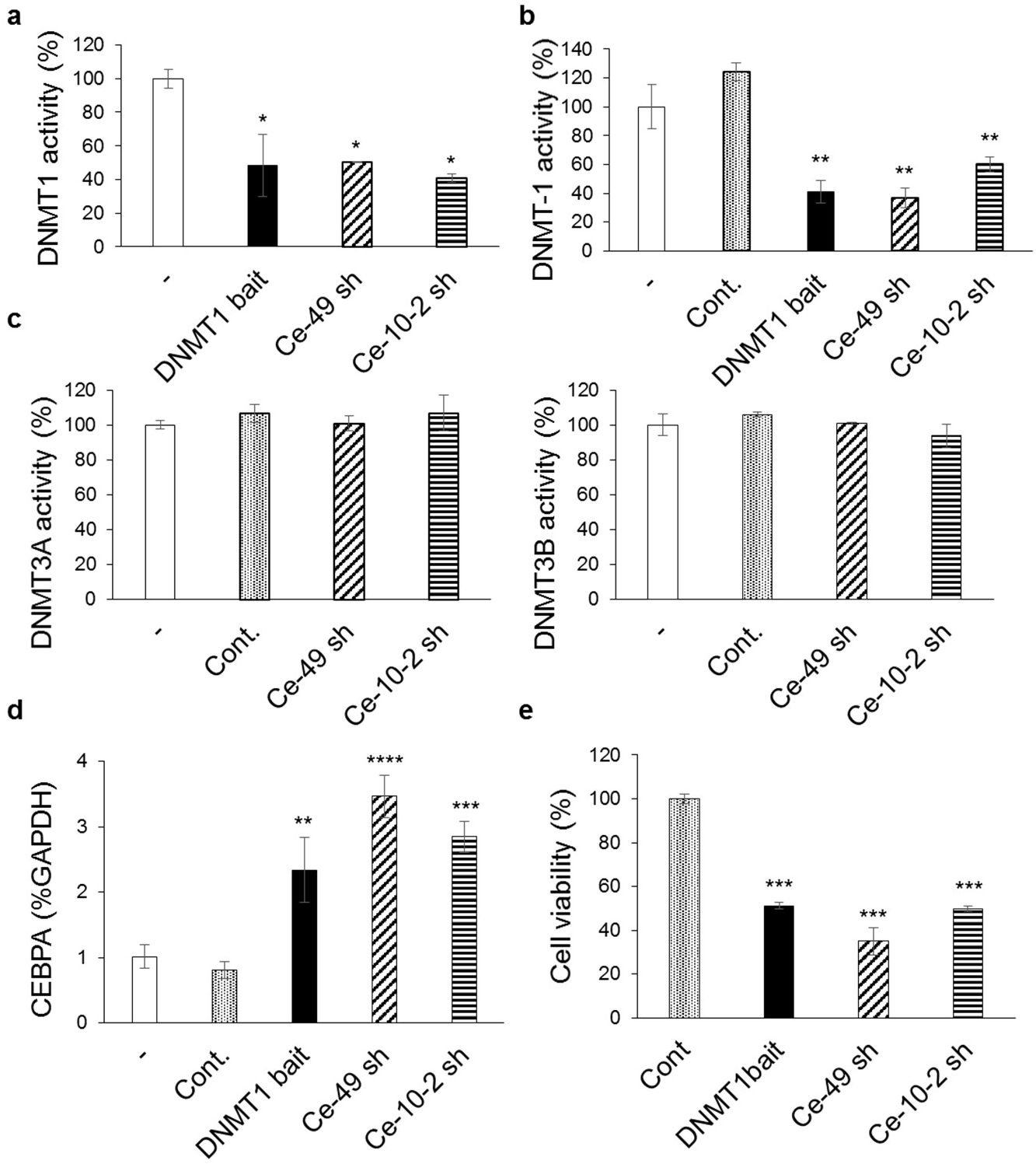
Functional activity of the best aptamers. (**a**) Activity of purified DNMT1 protein was analysed *in vitro* by DNMT1 inhibitor screening assay in the absence (-) or in the presence of indicated aptamers or DNMT1 bait. Results are expressed as percentage relative to the activity of DNMT1 protein alone. Mean ± SD is reported (n=2). Statistic by one-way ANOVA: *, p<0.05. (**b**) DNMT1 inhibitor screening assay was performed with nuclear extracts from K562 cells left untreated (-) or transfected with DNMT1 bait, indicated aptamers or mutR5 used as a control (Cont.). Results are expressed as percentage relative to the activity detected with untreated sample. Mean ± SD is reported (n=2). Statistic by one-way ANOVA (*versus* Cont.): **, p<0.01. (**c**) Activity of purified DNMT3A (*left*) or B (*right*) proteins was analysed in vitro in the absence (-) or in the presence of indicated aptamers and expressed as percentage relative to the activity of DNMT protein alone. Bars depict mean ± SD (n=2). (**d**) Levels of CEBPA were analysed by RT-qPCR in K562 cells left untreated (-) or transfected with indicated aptamers or Cont. for 72 hours. Mean ± SD is reported (n=3). Statistic by one-way ANOVA (*versus* Cont.): **, p<0.01; ***, p<0.001; ****, p<0.0001. (**e**) Cell viability of CML K562 cells transfected with indicated aptamers or Cont. for 72 hours. Bars depict mean ± SD (n=2). Statistic by one-way ANOVA (*versus* Cont.): ***, p<0.001.

Finally, we evaluated the transcriptional activation brought about by DNMT1 aptamers in K562 in which the *CEBPA* promoter is methylated and the mRNA is expressed at low-to-undetectable levels ^15^. Upon transfection with the DNMT1 aptamers, we observed effective increase of *CEBPA* levels (**Figure 7d**), whereas no changes occurred when the control oligonucleotide was used. As a result of DNMT1 inhibition and *CEBPA* upregulation^17,18^, K562 cell viability was reduced 60 to 50% upon aptamer transfection (**Figure 7e**) when compared to the non-targeting control and similar effect followed in Calu-1 and A549, non-small cell lung cancer cell lines (**Supplementary Figure 9**).

To estimate the extent of the effect on DNA methylation resulting from the DNMT1-specific aptamers, the genome-scale methylome profile was assessed by the EPIC array platform on K562 cells transfected with Ce-49 sh and Ce-10-2 sh. The differential methylation analyses revealed significant reduction of DNA methylation across thousands of CpG covered by the array in the aptamer-treated cells as compared to the control (**Figure 8a**). Nearly 16.000 and 14.000 differentially methylated regions (DMRs) were detected for the Ce-49 sh and Ce-10-2 sh samples, respectively (**Figure 8b** and **c**) with more than 60% overlap between the two (**Figure 8c**). Gene ontology (GO) analyses for “biological process” of genes corresponding to the overlapping hypo-methylated CpGs included among the top ranked, GO terms belonging to epigenetic modification, regulation of transcription and gene expression, consistently with the aptamer function (**Figure 8d**).

**Figure 8.**
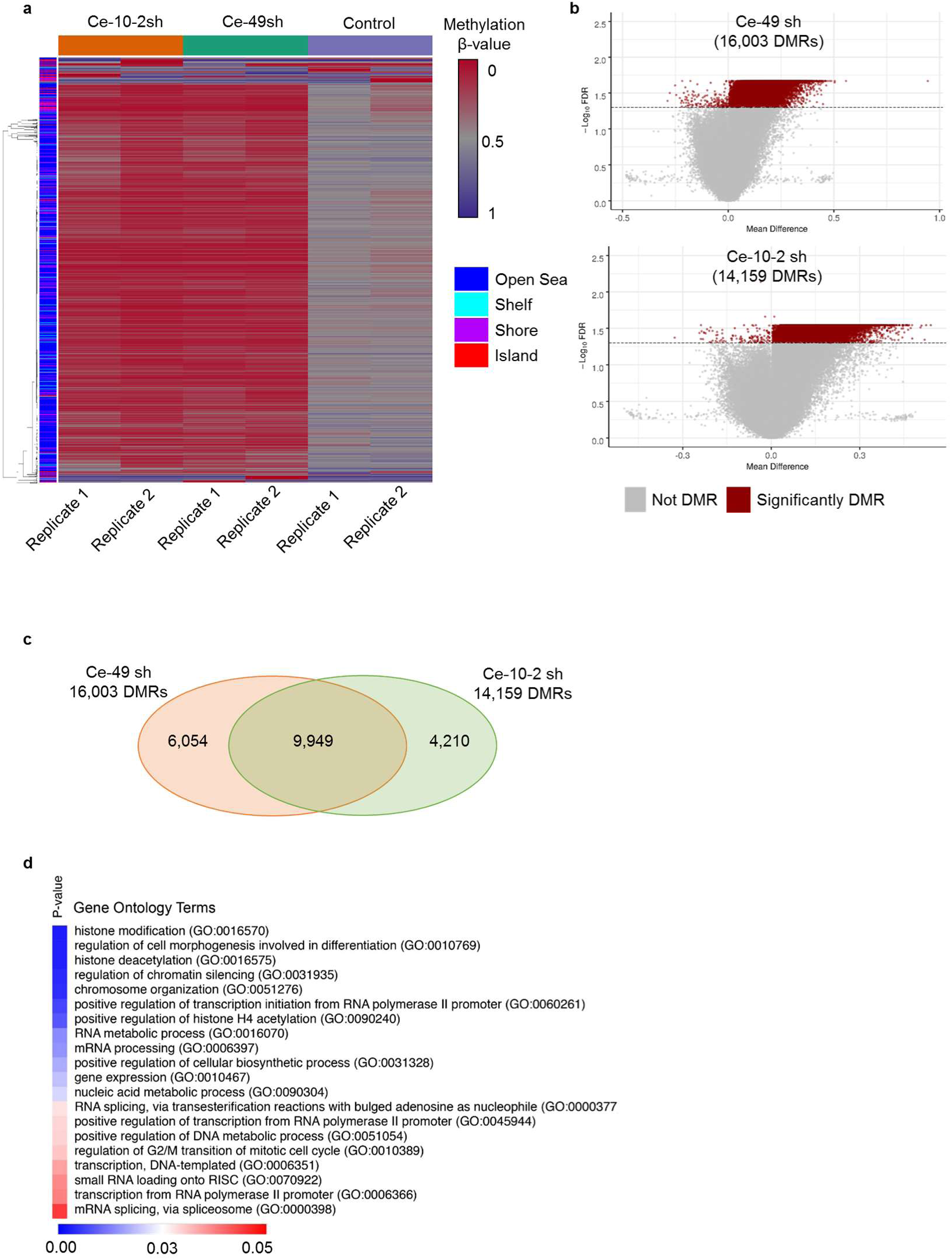
DNA methylation analyses. (**a**) Heatmap of differentially methylated CpG regions (DMR) in K562 transfected with Ce-49 sh, Ce-10-2 sh or or control (Cont.) aptamers. (**b**) Volcano plots reporting the significantly DMRs for Ce-49 sh (*upper panel*) and Ce-10-2 sh (*lower panel*). (**c**) DMRs overlapping between Ce-49 sh and Ce-10-2 sh. (**d**) Heatmap of top ranked Gene ontology terms for genes corresponding to the overlapping hypo-methylated CpGs shared by Ce-49 sh and Ce-10-2 sh.

Altogether, our results support the potential use of these newly generated aptamers as a novel approach to block DNMT1 activity, restore the expression of genes silenced by DNA methylation and reduce cell viability of cancer cells.

## Discussion

The study herein presented aims at neutralizing the major epigenetic player, DNMT1, using a newly developed RNA aptamer platform – the aptaDiR. On the assumption that RNAs are epigenetic modulators capable of inhibiting the activity of DNMT1^15,19,20^, we examined the possibility to merge this novel RNA feature to the key properties aptamers retain as ligands: the strong affinity and high specificity to a designated target.

We adopted an innovative doped SELEX strategy to isolate high-affinity DNMT1-specific RNA aptamers (**Figure 1**). Put simply, for the initial pool of selection we exploit the propensity of DNMT1 to bind stem-loop RNA^15^ and designed and mixed together three small libraries containing each a small degenerated region in the context of a short DiR sequence (R5) (**Figure 2**). This approach allowed the rapid evolution (only three rounds of selection) of ligands exhibiting stronger affinity and higher stability than the endogenous DNMT1-RNA bait (R5). As such, this strategy might have a broader range of application to generate aptamers targeting proteins binding nucleic acid and specifically-structured motifs present in larger transcripts as non-canonical RNAs.

Among the potential candidates, we shortlisted the most selective (with a KD of ∼ 70 nM) aptaDiRs (**Figure 3** and **4**). The aptaDiRs are unable to bind and inhibit the other two major DNMT family members: DNMT3A/3B, nor to bind to the unrelated proteins KAT5 and HSA (**Figure 5**). These molecules, stably form *in silico* and *in vitro* complexes with DNMT1 and inhibit DNMT1 activity reducing cell viability as expected for DNMT inhibitors (**Figure 7** and **8**).

Currently approved hypomethylating agents, such as the cytosine nucleoside analogues, lack selectivity and are characterized by low stability and high toxicity ^21^. In light of this, the herein aptaDiRs represent an alternative strategy to overcome the clinical limitations of the prevailing administered DNMT inhibitors.

Recently, Wang et al. ^22^ have reported the generation of aptamers targeting DNMT1 using as a starting pool a high complexity DNA library. However, it remains counterintuitive to opt for a single stranded (ss) DNA aptamer when DNMT1 has been reported to show stronger affinity to RNA than DNA ^15^. In fact, the DNMT1 aptamer emerging from their selection (Apt. #9) showed an affinity for DNMT1 of 0.77 μM, a value considerably higher than those detected for all the RNA sequences highlighted in our current and previous study ^15^. Our selection strategy is the first-of-its-kind as it preserves the secondary structure constraints required for DNMT1-RNA interaction while evolving aptamers with enhanced binding and stability for clinical translational purposes. RNA molecules are indeed more flexible and easier to manipulate to achieve stability as compared to ssDNAs^11^. The aptaDiR herein reported have affinity values in the nanomolar range, thus 10 times lower than the one indicated for the ssDNA aptamer by Wang et al. Importantly, the stability of the RNA molecules, which display an half-life longer than 48 hours in 85% human serum, considerably exceeds that of the reported Apt #9 with an half-life of 11 hours in 10% serum media ^22^.

In summary, this study presented two distinct novelties. First, it describes an original selection strategy that translates the DNMT1 neutralizing properties of specialized endogenous RNAs (DiRs) into an aptamer-based platform leading to the aptaDiRs. Second, it provides the first RNA-based approach that is highly selective, chemically stable and molecularly versatile to control DNA methylation for therapeutic use. Undoubtedly, the aptaDiRs are much more attractive for clinical applications than other nucleic acid-based compounds lacking these essential features. Furthermore, the ability of stem–loop structured aptamers to enter cells and localize within the nucleus has been demonstrated ^23,24^ and multiple strategies for an active delivery into the nucleus are now available ^25,26^. This is a critical point for the successful outcome of future *in vivo* applications and the translational feasibility of the aptaDiR platform.

Taken together these results provide a proof-of-concept for the generation of innovative, designated and cost-effective RNA-based epigenetic therapy. This strategy is not purely restricted to DNMT1 but conveyable to other epigenetic factors thereby broadening the clinical applications of RNA aptamers to multiple types of cancers or diseases associated with epigenetic alterations.

## Methods

### Library preparation and in vitro transcription

Randomized sub-libraries were purchased from Genomics and PCR amplified by using 0.05 U/μl Fire Pol DNA polymerase (Microtech) in a mix containing: 0.4 μM primers, 0.2 mM dNTPs. After 3 minutes initial denaturation at 95 °C, the protocol used was: 10 cycles of : 95 °C for 30 seconds, 64°C for 1 minute, 72 °C for 30 seconds; following final extension of 5 minutes at 72°C. Primers used were: forward (with T-7 RNA Polymerase promoter): 5’ TAATACGACTCACTATAGGGCTGAAGGGGTTACTGGG-3’; reverse: 5’-CTCCTCCCCGGGGCAGATA-3’. Equal amounts of each sub-libraries were mixed and transcribed. In vitro transcription was performed in the presence of 1 mM 2’-F Py using a mutant form of T7 RNA polymerase (Y639F). DNA template was incubated at 37°C overnight in a transcription mix containing: transcription buffer 1X (Epicentre Biotechnologies), 1 mM 2’F-Py (2’F-2’-dCTP and 2’F-2’-dUTP, TriLink Biotechnologies), 1 mM ATP, 1 mM GTP (Thermo scientific), 10 mM dithiothreitol (DTT) (Thermo scientific), 0.5 u/μl RNAse inhibitors (Roche), 5 μg/ml inorganic pyrophosphatase (Roche), and 1.5 u/μl of the mutant T7 RNA polymerase (T7 R&DNA polymerase, Epicentre Biotechnologies). After transcription, any leftover DNAs were removed by DNase I (Roche) digestion and RNAs were recovered by phenol:chloroform extraction, ethanol precipitation and gel purification on a denaturing 8% acrylamide/7 M Urea gel ^27^.

*In vitro* transcription in the presence of a mixture of 5’-biotin-G-monophosphate (TriLink Biotechnologies) and GTP (molar ratio 3:2) were performed for the biotinylation of the long aptamers.

### SELEX procedure

For the SELEX strategy, GST-tagged DNMT1 was used as target for the selection. Recombinant Human DNMT1 with an N-terminal GST tag was purchased from Active Motif and Pierce Glutathione Magnetic Beads (Thermo Scientific) were used to separate aptamer-GST-tagged protein complexes.

Before each cycle of SELEX, the 2’F-Py RNA pool was dissolved in RNAse free water and subjected to denaturation/renaturation steps of 85 °C for 5 min, ice for 2 min and 37°C for 5 min. The RNA-protein incubation was performed in Binding Buffer (BB: 5 mM TrisHCl pH 7,5; 5 mM MgCl2; 1 mM DTT; 100 mM NaCl). At each cycle, the pool was first incubated for 30 min with Glutathione Magnetic Beads with a gentle rotation, as counter-selection step, and then the unbound RNA was recovered on a magnetic separator and used for selection. The recovered sequences were incubated with GST-tagged DNMT1 at room temperature for 30 min with a gentle rotation. Aptamer-protein complexes were purified on magnetic beads. The unbound was removed on a magnetic separator and the beads containing RNA-protein complexes were washed with BB. Bound RNAs were recovered by TriFast (Euroclone) extraction and RT-PCR and finally transcribed for the following round.

### RT-PCR

RNA recovered at each SELEX round was reverse transcribed by using M-MuLV reverse Transcriptase (Roche) in a mix containing a specific 5X buffer (50 mM Tris-HCl pH 8.3, 40 mM KCl, 6 mM MgCl2, 10mM DTT), 0.8 µM of reverse primer, 1mM dNTPs. The protocol used for the reaction was: 30 min a t 42°C and 30 min at 50°C. The obtained product was then amplified by error prone PCR reaction in the presence of high MgCl2 (7.5 mM) and dNTPs (1 mM).

### Cloning, sequencing Bioinformatics analyses

The final pool from SELEX was amplified by PCR, including in the program a 15 minutes final extension at 72°C to introduce A-overhangs. Individual sequences were cloned with TOPO-TA Cloning Kit (Invitrogen) according to the manufacturer’ s instruction. Single white clones were grown and DNA was extracted with plasmid Miniprep kit (Qiagen) and sequenced by Eurofins Genomics. Single aptamer sequences obtained were analysed by using Multiple *Muscle* Algorithm. Aptamer secondary structures were predicted by using *RNAstructure* software.

### Synthetic oligonucleotides

Short RNAs were purchased from Trilink Biotechnologies with 2’-FPy all along the sequences. Biotin was added at the 3’-end of the sequences. Before each analysis, aptamers were kept 5 min at 85 °C, 2 min on ice and 5 min at 37 °C to enable the folding into their active conformations.

### ELONA Assay

Microtiter High Binding plate (Nunc MaxiSorp) wells were coated with 30 nM of His-tagged DNMT1 (Active Motif) or HSA overnight at 4°C. All subsequent steps were performed at room temperature. The plate wells were washed once with PBS and then blocked with 300 µl 3% BSA (AppliChem) in PBS for 2 hours. After two washes, biotinylated aptamers dissolved in PBS (100 µL) were added for 2 hours. After three washes, streptavidin-conjugated horseradish peroxidase (HRP) (Sigma Aldrich) was incubated (1:10000 dilution) for 1 hour. Then, the plate was washed four times and the 3,3′,5,5′-Tetramethylbenzidine (TMB) substrate solution was added. The reaction was stopped with sulfuric acid (H2SO4) 0.16 M, forming a yellow reaction product. Signal intensity was measured with microplate Reader (Thermo Scientific) at 450 nm.

### Human Serum Stability assay

Oligonucleotides were incubated at 4 μM concentration in 85% human serum from T0 to 7 days. Type AB Human Serum provided by Sigma Aldrich was used. At each time point, RNAs (4 μl, 16 pmoles) were recovered and incubated for 1 hours at 37 °C with 2 μl of proteinase K solution (20 mg/ml) in order to remove serum proteins that interfere with RNA electrophoretic migration. Following proteinase K treatment, 12 μl of denaturing dye (95% formamide, 10 mM EDTA, Bromophenol Blue) was added to samples that were then stored at −80 °C. All time point samples were separated by electrophoresis into 15% acrylamide/7 M Urea gel. The gel was stained with ethidium bromide and visualized by UV exposure.

### Bio-Layer Interferometry technology system (BLItz)

Bio-Layer Interferometry measurements were performed using a BLItz system and AR2G tips (Amine Reactive biosensors of 2nd Generation) (ForteBio Inc). After pre-hydration for 10 min in PBS buffer, the AR2G tips were efficiently functionalized with DNMT1 (Active Motif), DNMT3A (Abcam), DNMT3B (Abcam) and KAT 5 (KAT5-1350H; Creative Biomart) proteins following the manufacturer’s instructions. The AR2G tips enable the coupling of proteins to carboxylate groups on the biosensor surface via accessible amine groups. Accordingly, tips were activated with an EDC (0.2 M): NHS (0.05 M) coupling mixture for 300 sec, then each ligand at the concentration of 20 μg/mL diluted in 100 mM MES pH 5.0 was exposed for 180 sec to distinct biosensors. The average immobilization levels ranged between 2.0 and 2.5 nm). Unused activated carboxylated groups on the tips surface were reacted with 1.0 M ethanolamine hydrochloride, pH 8.5 for 120 sec. After activation, a regeneration step with 10 mM NaOH was performed to minimize non-specific binding. Blank biosensors were similarly prepared by activation and deactivation steps and were used to collect and subtract reference interferograms. For dose-dependent assays increasing concentrations of Ce-49 sh (concentrations ranging between 1.0 nM and 750 nM), Ce-10-2 sh (concentrations ranging between 10 nM and 750 nM) or DNMT1 bait (concentrations ranging between 100 nM and 2.0 μM) were used. Each run was performed following the reported steps: initial baseline (30 sec), association (120 sec) and dissociation (120 sec), regeneration (2 x 30 sec, 10 mM NaOH). The association step was performed using 4.0 μL of ligand solution placed in a drop holder drop, setting the shaker speed at 2000 rpm, according to the manufacturer’s instructions. Duplicate or triplicate experiments were performed at increasing aptamer concentrations. Reference interferograms were subtracted from experimental values before data processing to reduce the background. Data were exported from the BLItz Pro 1.2 software and re-plotted with GraphPad Software. Plateau values of binding as reflected by changes in optical thickness (nm) at 120 sec were used to calculate the affinity constant (KD) by applying a non-linear curve fitting and one binding site hyperbola as model (GraphPad Prism).

### REMSA

Aptamers (15 pmol) and DNA double stranded oligonucleotides were end-labelled with [γ-^32^P] ATP (Perkin Elmer) and T4 polynucleotide kinase (New England Biolabs). Reactions were incubated at 37 °C for 1 h and then passed through G-25 spin columns (GE Healthcare) according to the manufacturer’s instructions to remove unincorporated radioactivity. Labelled samples were gel purified on 10% polyacrylamide gels prepared with 0.5x TBE buffer. Binding reactions were carried out in 10-μl volumes in the following buffer: 5mM Tris, pH 7.4, 5mM MgCl2, 1mM dithiothreitol (DTT), 3% (v/v) glycerol, 100mM NaCl. 0.8μM of purified DNMT1 protein (Active Motif) was incubated with 30000 cpm of ^32^P-labelled aptamers and single or double-stranded, RNA or DNAs, respectively. All reactions were assembled on ice and incubated for 30 minutes at room temperature. Samples were separated on 6% native polyacrylamide gels (0.53 Tris/Borate/EDTA (TBE); 4 °C; 2.5 hs at 190V). Gels were fixed, dried and exposed to X-ray film.

### Aptamer-mediated pulldown

K562 cells were lysed with 10 mM Tris-HCl pH 7.5 containing 200 nM NaCl, 5 mM ethylenediaminetetraacetate (EDTA), 0.1% Triton X-100 and protease inhibitors. Extracts (500 μg in 0.5 ml lysis buffer) were incubated for 30 min with 200 nM heat denatured biotinylated aptamers with rotation. Following three washings with PBS cells, aptamer-protein complexes were purified on streptavidin beads (Thermo Fisher Scientific) for 2 hours. Beads were washed three times with PBS and bound proteins were recovered by adding Laemmli buffer and then analysed by immunoblotting with anti-DNMT1 antibody (Active Motif).

### Modelling and Molecular Dynamics Simulations

The three-dimensional structure of the human DNMT1 protein was created by homology modelling with MODELLER 9.22 program using the mouse DNMT1 in complex with its DNA substrate solved structure as template, downloaded from the Protein Data Bank database (PDB: 4d4a) ^28^. Three-dimensional models of R5, mutR5, Ce-49 sh and Ce10-2 sh RNA aptamers were obtained employing the MC-Fold-MC-Sym pipeline ^29^ and optimized with 100ns of MD simulations in waters. Aptamer representative conformations were extracted from each MD trajectories by clustering method and subsequently used to build the relative complex with the human DNMT1 modelled protein. In details, the complexes were built by manually docking ^30^ each aptamers conformation into the catalytic region of the human DNMT1 structure, using the DNA substrate coordinates as solved in the 4d4a pdb, as guide to define the binding pose. The initial complexes of human DNMT1 with R5, Ce-49 sh and Ce10-2 sh aptamers were then subjected to MD simulations. MD simulations were run with GROMACS 5.0.5 ^31^ all the systems were solvated in an octahedron box using the TIP3P water models ^32^ with a 1.1 nm distance to the border of the molecule, simulating standard biological conditions by considering a 150 mM KCl concentration and additional ions to neutralize. Electrostatic interactions were treated using the particle mesh Ewald method and Berendsen algorithm to control temperature and pressure ^33^, following the indications dictated by the ABC consortium (https://bisi.ibcp.fr/ABC/Protocol.html) and previous protocols ^34-38^. In all the systems, waters were firstly relaxed by energy minimization and 10 ps of simulations at 300 K, restraining the protein and RNA atomic positions with a harmonic potential. Then, the systems were heated up gradually to 300 K in a six step phases starting from 50 K, finally the simulations were run in NPT standard conditions for 300 ns without restraints. GROMACS^31^, VMD^39^ and Pymol ^40^ packages were used to analyse all the MD trajectories. Clustering analyses of the last half time of each MD simulation were performed to extract representative conformations using the gromos clustering method with the algorithm described by Dura et al. ^41^. For each cluster, the structure exhibiting the lowest RMSD relative to all the other members of the cluster was selected as representative.

### Cells and transfection

K562 cells, A549 and Calu-1 NSCLC cells were grown in RPMI medium supplemented with 10% FBS (Sigma). Transfections were performed using serum-free Opti-MEM and Lipofectamine 2000 reagent (Thermo Fisher Scientific) according to the manufacturer’s protocol. Cells were transfected with 100 nM of RNAs previously subjected to denaturation/renaturation steps.

### Quantitative RT-PCR (RT-qPCR)

To analyze gene mRNA level, 1 μg of total RNA was reverse transcribed with iScript cDNA Synthesis Kit and amplified by real-time quantitative PCR with IQ-SYBR Green supermix (Bio-Rad, Hercules, CA, USA). ΔΔCt method was used for relative mRNA quantization by applying the equation 2^-ΔΔCt^.

Primers used were: human *CEBPA*: Forward: 5’CCGCTCCTCCACGCCTGTCCTTAG-3’; Reverse: 5’-GCCCCACAGCCAGATCTCTAGGTC-3’; GAPDH (housekeeping control): Forward: 5’-CTTTGTCAAGCTCATTTCCTGG-3’; Reverse: 5’-TCTTCCTCTTGTGCTCTTGC-3’.

### Cell viability assay

Cells were seeded in 96-well plates (3 × 10^3^ cells/well) and were transfected with indicated sequences (100 nM). Following 72 hours, cell viability was assessed by CellTiter 96 Proliferation Assay (Promega).

### DNA methylation analysis

Genomic DNAs from K562 cells transfected with aptamers or control (mutR5) were extracted with DNeasy Blood & Tissue Kit (Qiagen). Samples were analysed by Diagenode EPIC methylation array after bisulfite conversion.

The methylation array data utilized for this manuscript can be accessible through GEO with accession GSE154471

The EPIC methylation array was analysed using the Bioconductor package RnBeads (v2.4.0)^42,43^ using the hg19 annotations. Background normalization method was set to “enmix.oob” with “swan” used for normalization. When setting up the sva covariates, only the “aptamers” designation was used. Differential methylation analysis was then performed comparing the respective “control” and “treatment” replicates. The heatmap output from RnBeads was used with minor adjustments for publication. For volcano plots, EnhancedVolcano (v1.4.0)^44^ was used. CpG sites with an adjusted FDR < 0.05 were set to dark red for plotting. CpG sites were annotated to their closest gene using the variable annotations provided with the array platform using a custom R script. For Gene Onology analysis, hypo-methylated CpG sites (with mean.diff >=0 value, FDR < 0.05) were selected, their HGCN gene ids extracted and analysed using Enrichr^45,46^. The results reported herein correspond to the resulting “GO Biological Process” results.

### Statistics

The statistical evaluation, we applied student’*t*-test for caparison between two groups (Figure 2a and Figure 3) and one-way ANOVA with multiple comparisons for functional assays (Figure 7 and Supplementary 9). GraphPad Prism Software was used. Values of p≤ 0.05 were considered statistically significant.

## Data and software availability

Data are available on the gene omnibus database under the accession ID number: GSE154471 (https://www.ncbi.nlm.nih.gov/geo/query/acc.cgi?acc=GSE154471)

## Acknowledgements

This work was supported by the National Cancer Institute of the National Institutes of Health under Award Number R00CA188595; the Italian Association for Cancer Research (AIRC) Start-Up grant #2014-15347, the Fondazione Cariplo N. 2016-0476 and the Giovanni Armenise-Harvard Foundation to ADR; the Italian Ministry of Economy and Finance to the CNR for the Project FaReBio di Qualità and the Asian Fund for Cancer Research Limited to VDF; the Italian Ministry of Health, GR-2011-02352546 to CLE; The Italian Ministry for University and Research (MUR) for the project PRIN n° 2015783N45 and project NEON, n° ARS01_00769 to MR; the Regione Campania for the projects NANOCAN and SATIN to MR. MB was supported by a PhD Fellowship from the Italian Ministry of Education, Universities, and Research to the University of Eastern Piedmont. We wish to thank Prof. Daniel Tenen for constructive discussion and insightful suggestions.

## Author Contributions

ADR, CLE, and VDF conceived and designed the study and wrote the manuscript; ADR; CLE; IA; SA; SU; MB; SC performed experiments; MAB performed bioinformatics and statistical analyses; MR, AKE provided intellectual supports and revised the manuscript.

## Competing interests

Authors declare no competing interest.

## Notes

https://www.ncbi.nlm.nih.gov/geo/query/acc.cgi?acc=GSE154471

### Competing Interest Statement

The authors have declared no competing interest.

## References

1 Li, E., Beard, C. & Jaenisch, R. Role for DNA methylation in genomic imprinting. Nature 366, 362–365, doi:10.1038/366362a0 (1993).

2 Razin, A. & Riggs, A. D. DNA methylation and gene function. Science 210, 604–610, doi:10.1126/science.6254144 (1980).

3 Ehrlich, M. et al. Amount and distribution of 5-methylcytosine in human DNA from different types of tissues of cells. Nucleic Acids Res 10, 2709–2721, doi:10.1093/nar/10.8.2709 (1982).

4 Goll, M. G. & Bestor, T. H. Eukaryotic cytosine methyltransferases. Annu Rev Biochem 74, 481–514, doi:10.1146/annurev.biochem.74.010904.153721 (2005).

5 Mair, B., Kubicek, S. & Nijman, S. M. Exploiting epigenetic vulnerabilities for cancer therapeutics. Trends Pharmacol Sci 35, 136–145, doi:10.1016/j.tips.2014.01.001 (2014).

6 Feinberg, A. P. & Tycko, B. The history of cancer epigenetics. Nat Rev Cancer 4, 143–153, doi:10.1038/nrc1279 (2004).

7 Zaman, A. & Bivona, T. G. Emerging application of genomics-guided therapeutics in personalized lung cancer treatment. Ann Transl Med 6, 160, doi:10.21037/atm.2018.05.02 (2018).

8 Santi, D. V., Norment, A. & Garrett, C. E. Covalent bond formation between a DNA-cytosine methyltransferase and DNA containing 5-azacytosine. Proc Natl Acad Sci U S A 81, 6993–6997, doi:10.1073/pnas.81.22.6993 (1984).

9 Lyko, F. & Brown, R. DNA methyltransferase inhibitors and the development of epigenetic cancer therapies. J Natl Cancer Inst 97, 1498–1506, doi:10.1093/jnci/dji311 (2005).

10 Yin, Y. et al. Impact of cytosine methylation on DNA binding specificities of human transcription factors. Science 356, doi:10.1126/science.aaj2239 (2017).

11 Keefe, A. D., Pai, S. & Ellington, A. Aptamers as therapeutics. Nat Rev Drug Discov 9, 537–550, doi:10.1038/nrd3141 (2010).

12 Mercier, M. C., Dontenwill, M. & Choulier, L. Selection of Nucleic Acid Aptamers Targeting Tumor Cell-Surface Protein Biomarkers. Cancers (Basel) 9, doi:10.3390/cancers9060069 (2017).

13 Zhang, Y., Lai, B. S. & Juhas, M. Recent Advances in Aptamer Discovery and Applications. Molecules 24, doi:10.3390/molecules24050941 (2019).

14 Zhu, G. & Chen, X. Aptamer-based targeted therapy. Adv Drug Deliv Rev 134, 65–78, doi:10.1016/j.addr.2018.08.005 (2018).

15 Di Ruscio, A. et al. DNMT1-interacting RNAs block gene-specific DNA methylation. Nature 503, 371–376, doi:10.1038/nature12598 (2013).

16 Abdiche, Y., Malashock, D., Pinkerton, A. & Pons, J. Determining kinetics and affinities of protein interactions using a parallel real-time label-free biosensor, the Octet. Anal Biochem 377, 209–217, doi:10.1016/j.ab.2008.03.035 (2008).

17 Brown, K. D. & Robertson, K. D. DNMT1 knockout delivers a strong blow to genome stability and cell viability. Nat Genet 39, 289–290, doi:10.1038/ng0307-289 (2007).

18 Lourenco, A. R. & Coffer, P. J. A tumor suppressor role for C/EBPalpha in solid tumors: more than fat and blood. Oncogene 36, 5221–5230, doi:10.1038/onc.2017.151 (2017).

19 Savell, K. E. et al. Extra-coding RNAs regulate neuronal DNA methylation dynamics. Nat Commun 7, 12091, doi:10.1038/ncomms12091 (2016).

20 Zhao, Y., Sun, H. & Wang, H. Long noncoding RNAs in DNA methylation: new players stepping into the old game. Cell Biosci 6, 45, doi:10.1186/s13578-016-0109-3 (2016).

21 Foulks, J. M. et al. Epigenetic drug discovery: targeting DNA methyltransferases. J Biomol Screen 17, 2–17, doi:10.1177/1087057111421212 (2012).

22 Wang, L. et al. A DNA aptamer for binding and inhibition of DNA methyltransferase 1. Nucleic Acids Res 47, 11527–11537, doi:10.1093/nar/gkz1083 (2019).

23 Trinh, T. L. et al. A Synthetic Aptamer-Drug Adduct for Targeted Liver Cancer Therapy. PLoS One 10, e0136673, doi:10.1371/journal.pone.0136673 (2015).

24 Xiang, Q. et al. Suppression of FOXM1 Transcriptional Activities via a Single-Stranded DNA Aptamer Generated by SELEX. Sci Rep 7, 45377, doi:10.1038/srep45377 (2017).

25 Kotula, J. W. et al. Aptamer-mediated delivery of splice-switching oligonucleotides to the nuclei of cancer cells. Nucleic Acid Ther 22, 187–195, doi:10.1089/nat.2012.0347 (2012).

26 Nag, O. K. & Delehanty, J. B. Active Cellular and Subcellular Targeting of Nanoparticles for Drug Delivery. Pharmaceutics 11, doi:10.3390/pharmaceutics11100543 (2019).

27 Catuogno, S., Esposito, C. L. & de Franciscis, V. Developing Aptamers by Cell-Based SELEX. Methods Mol Biol 1380, 33–46, doi:10.1007/978-1-4939-3197-2_3 (2016).

28 Song, J., Teplova, M., Ishibe-Murakami, S. & Patel, D. J. Structure-based mechanistic insights into DNMT1-mediated maintenance DNA methylation. Science 335, 709–712, doi:10.1126/science.1214453 (2012).

29 Parisien, M. & Major, F. The MC-Fold and MC-Sym pipeline infers RNA structure from sequence data. Nature 452, 51–55, doi:10.1038/nature06684 (2008).

30 Autiero, I., Langella, E. & Saviano, M. Insights into the mechanism of interaction between trehalose-conjugated beta-sheet breaker peptides and Abeta(1-42) fibrils by molecular dynamics simulations. Mol Biosyst 9, 2835–2841, doi:10.1039/c3mb70235a (2013).

31 Hess, B., Kutzner, C., van der Spoel, D. & Lindahl, E. GROMACS 4: Algorithms for Highly Efficient, Load-Balanced, and Scalable Molecular Simulation. J Chem Theory Comput 4, 435–447, doi:10.1021/ct700301q (2008).

32 Pekka, M. a. N. L. Structure and Dynamics of the TIP3P, SPC, and SPC/E Water Models at 298 K.. J. Phys. Chem. A 105, 7 (2001).

33 Berendsen, H. J. C. G.. J. R.; Straatsma, T. P. The missing term in effective pair potentials. J. Phys. Chem. 91, 3, doi:10.1021/j100308a038 (1987).

34 Piacenti, V. et al. A combined experimental and computational study on peptide nucleic acid (PNA) analogues of tumor suppressive miRNA-34a. Bioorg Chem 91, 103165, doi:10.1016/j.bioorg.2019.103165 (2019).

35 Autiero, I., Ruvo, M., Improta, R. & Vitagliano, L. The intrinsic flexibility of the aptamer targeting the ribosomal protein S8 is a key factor for the molecular recognition. Biochim Biophys Acta Gen Subj 1862, 1006–1016, doi:10.1016/j.bbagen.2018.01.014 (2018).

36 Chawla, M., Autiero, I., Oliva, R. & Cavallo, L. Energetics and dynamics of the non-natural fluorescent 4AP:DAP base pair. Phys Chem Chem Phys 20, 3699–3709, doi:10.1039/c7cp07400j (2018).

37 Roviello, G. N., Roviello, V., Autiero, I. & Saviano, M. Solid phase synthesis of TyrT, a thymine-tyrosine conjugate with poly(A) RNA-binding ability. RSC Adv 6, 27607–27613, doi:10.1039/c6ra00294c (2016).

38 Autiero, I., Saviano, M. & Langella, E. In silico investigation and targeting of amyloid beta oligomers of different size. Mol Biosyst 9, 2118–2124, doi:10.1039/c3mb70086k (2013).

39 Humphrey, W., Dalke, A. & Schulten, K. VMD: visual molecular dynamics. J Mol Graph 14, 33-38, 27-38, doi:10.1016/0263-7855(96)00018-5 (1996).

40 DeLano, W. L. PyMOL. DeLano Scientific, San Carlos, CA, 700. (2002).

41 Daura, X. et al. Peptide Folding: When Simulation Meets Experiment. Angewandte Chemie International Edition 38, 236–240, doi:10.1002/(sici)1521-3773(19990115)38:1/2<236::Aid-anie236>3.0.Co;2-m (1999).

42 Muller, F. et al. RnBeads 2.0: comprehensive analysis of DNA methylation data. Genome Biol 20, 55, doi:10.1186/s13059-019-1664-9 (2019).

43 Assenov, Y. et al. Comprehensive analysis of DNA methylation data with RnBeads. Nat Methods 11, 1138–1140, doi:10.1038/nmeth.3115 (2014).

44 Blighe K R. S., Lewis M EnhancedVolcano: Publication-ready volcano plots with enhanced colouring and labeling. R package version 1.6.0, < https://github.com/kevinblighe/EnhancedVolcano > (2020).

45 Chen, E. Y. et al. Enrichr: interactive and collaborative HTML5 gene list enrichment analysis tool. BMC Bioinformatics 14, 128, doi:10.1186/1471-2105-14-128 (2013).

46 Kuleshov, M. V. et al. Enrichr: a comprehensive gene set enrichment analysis web server 2016 update. Nucleic Acids Res 44, W90–97, doi:10.1093/nar/gkw377 (2016).

