## Supplementary data for "Targeted systematic evolution of an RNA platform neutralizing DNMT1 function and controlling DNA methylation"

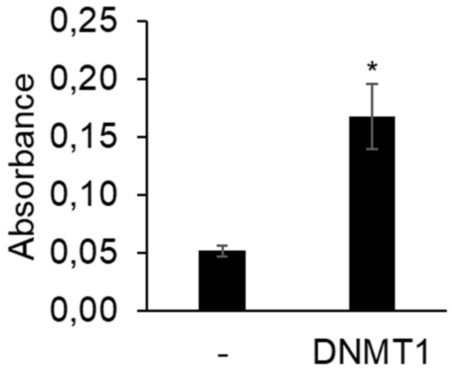


**Supplementary Figure 1. DNMT1 bait binding on DNMT1 purified protein.** Binding ability of R5 sequence modified with 2’-FPy (DNMT1 bait) on DNMT1 purified protein was detected by ELONA. Bars indicate mean ±SD (n=2). Statistics by *t*-test: *, $p<0.05$.

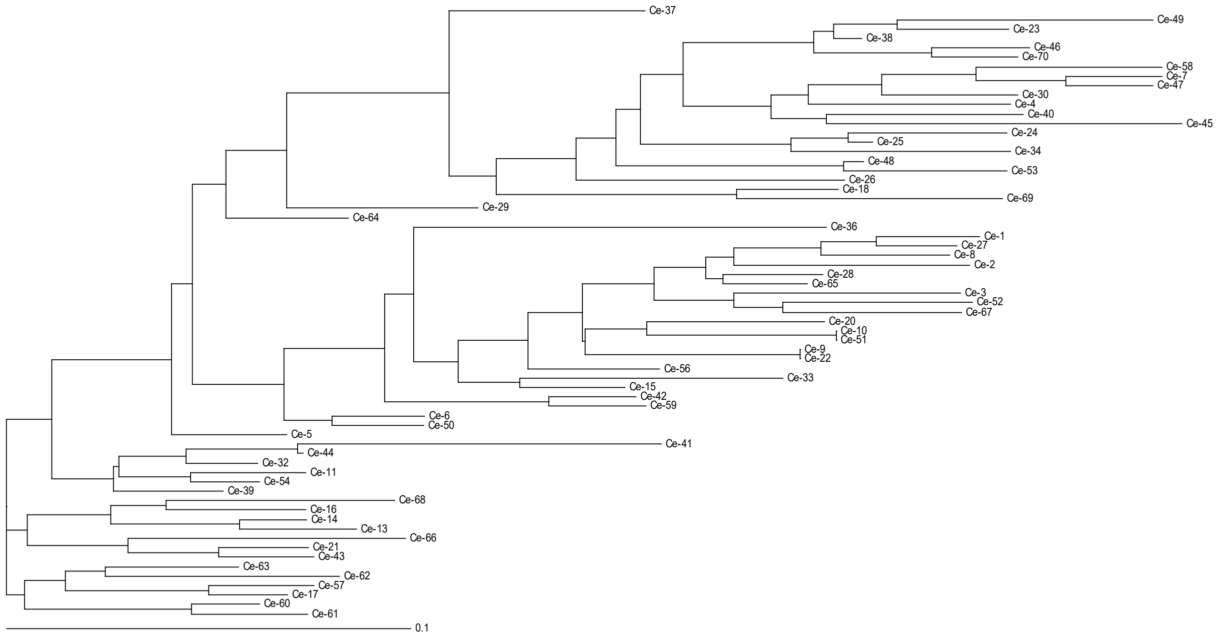


**Supplementary Figure 2. Families of individual aptamers from SELEX**. Dendrogram of the individual sequences cloned after the SELEX rounds. The three sequences chosen for further analyses are boxed.

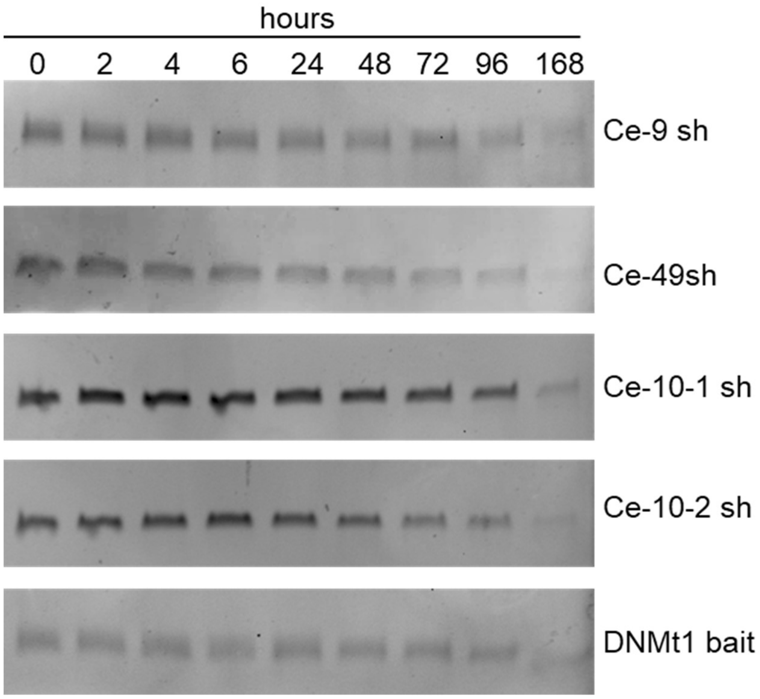


**Supplementary Figure 3. Serum stability of short individual aptamers**. Short aptamers and DNMT1 bait serum stability were measured in 85% human serum for indicated times. At each time point, RNA-serum samples were collected and evaluated by electrophoresis with 15% denaturing polyacrylamide gel. Gels were stained with ethidium bromide.

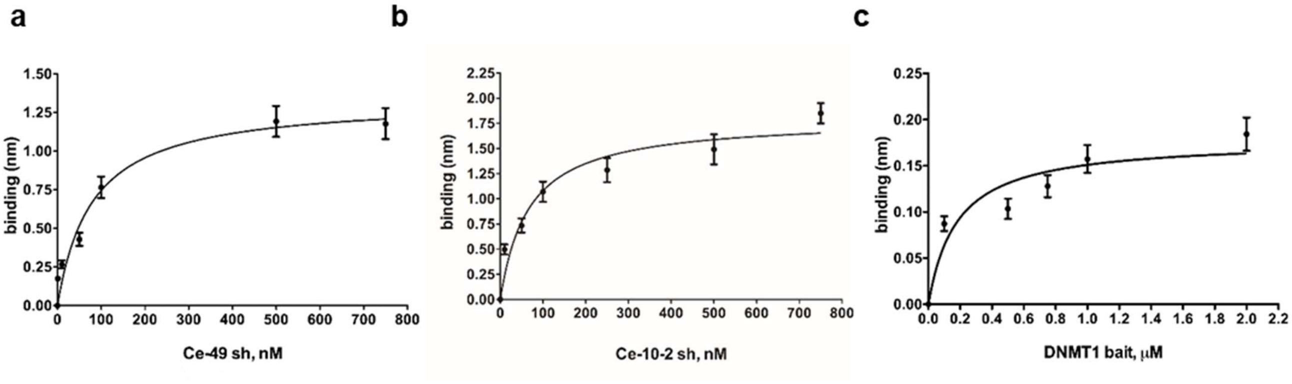


**Supplementary Figure 4. BLI affinity analyses**. Binding curves derived from BLItz analyses of Ce-49 sh (**a**), Ce-10-2 sh (**b**) or DNMT1 bait (**c**). Curves were fitted with a 1:1 binding model using GraphPad Prism 6.

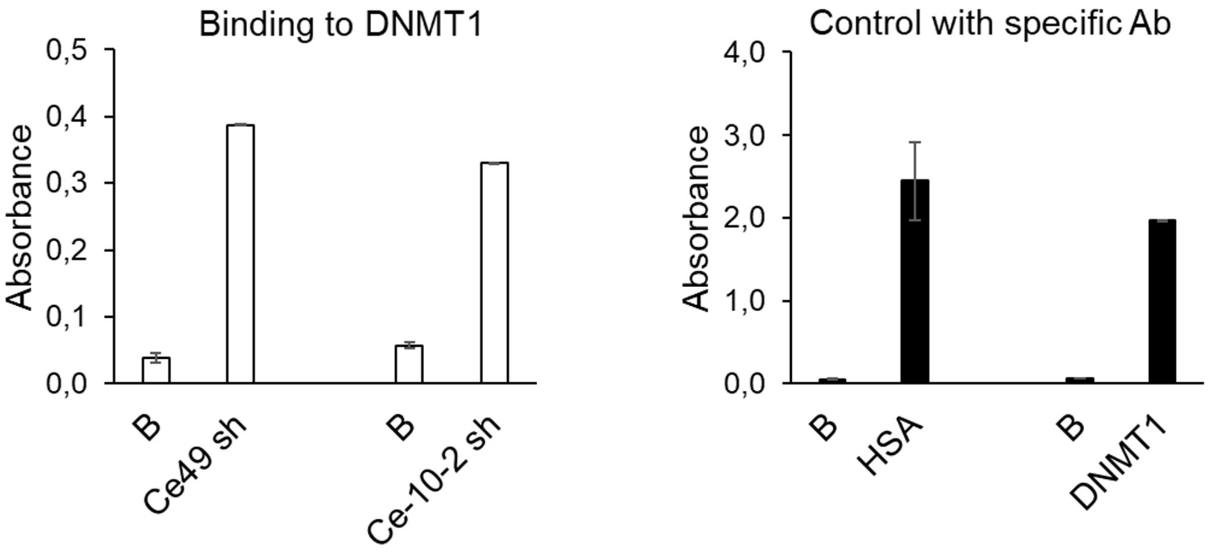


**Supplementary Figure 5. HSA assay controls.** For HSA assay, controls were performed by incubating aptamers at 200 nM with DNMT1 protein (*left*) or using specific antibodies to check the effective coating of the plates (*right*).

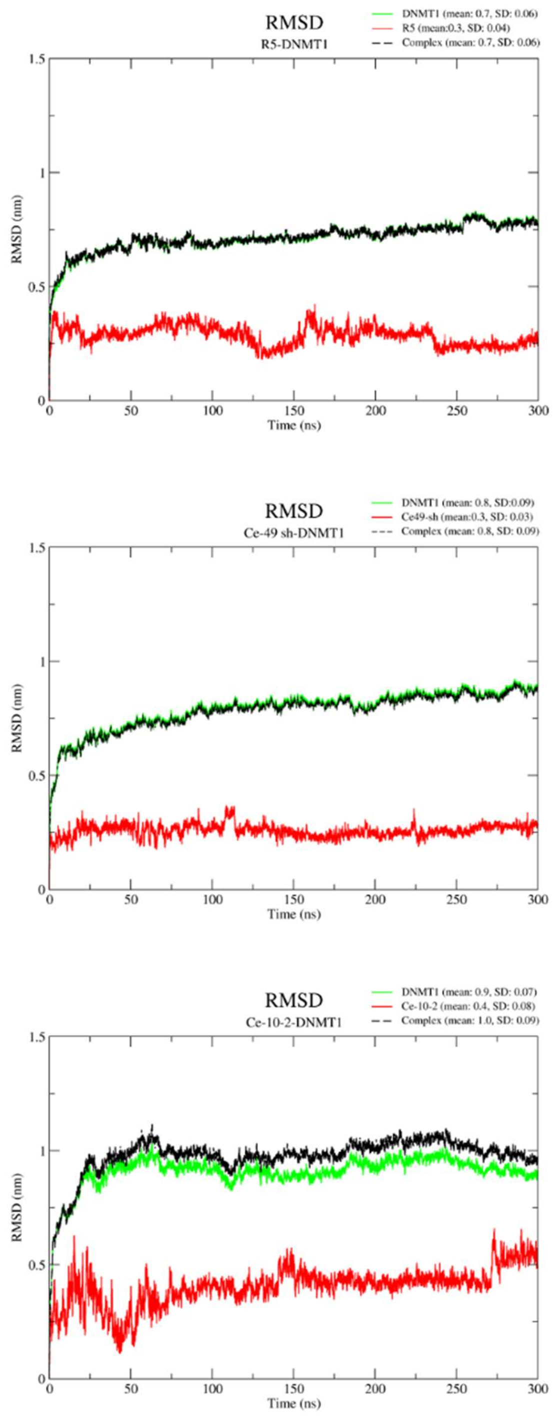


**Supplementary Figure 6. Time evolution of the RMSD values respect to the starting models.** The RMSD have been computed considering the C alpha and C5' atomos of the protein and RNA, respectively. The following colour code was used: overall complexes, black; green, protein residues; red, RNA residues.

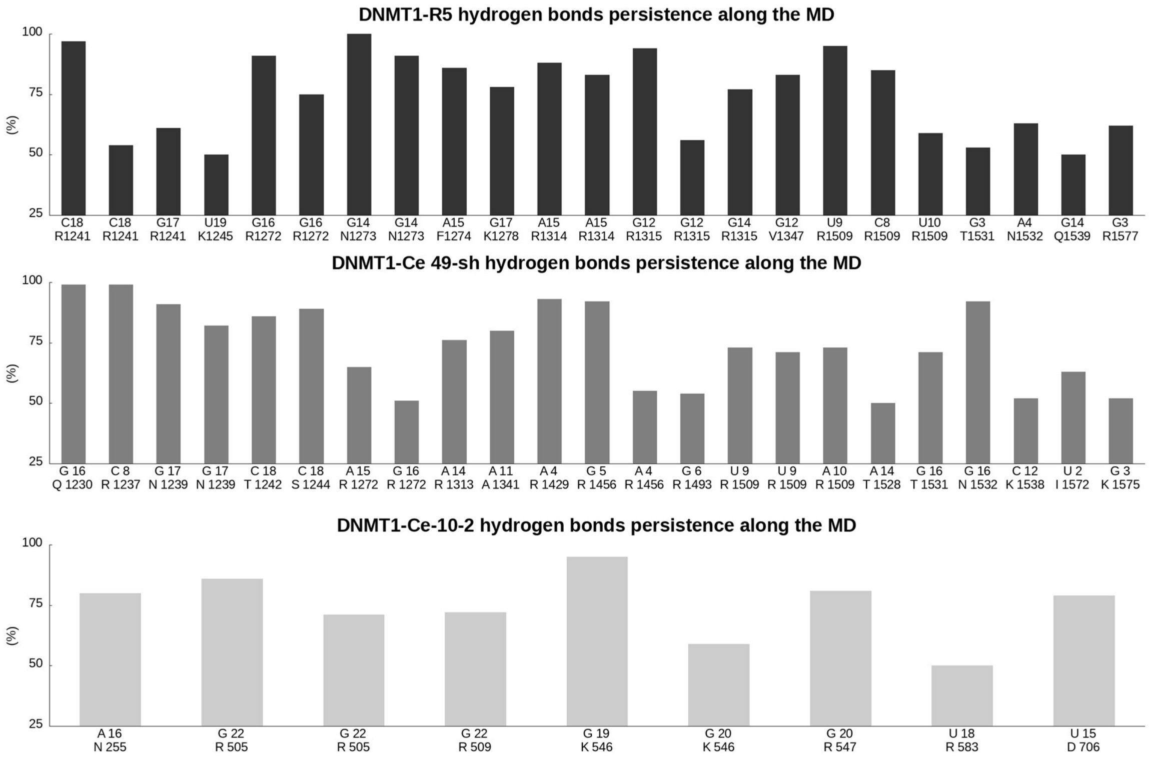


**Supplementary Figure 7. Hydrogen bonds at R5- or aptamers-DNMT1 interfaces**. Percentage of existence of hydrogen bonds at R5- or aptamers-DNMT1 interfaces, during the last 150 ns of simulation time, of R5-DNMT1: top panel, Ce-49 sh-DNMT1: central panel, and Ce-10-2 sh-DNMT1: bottom panel. The couple of protein-nucleobase residues involved into each hydrogen bond are indicated at the x axis.

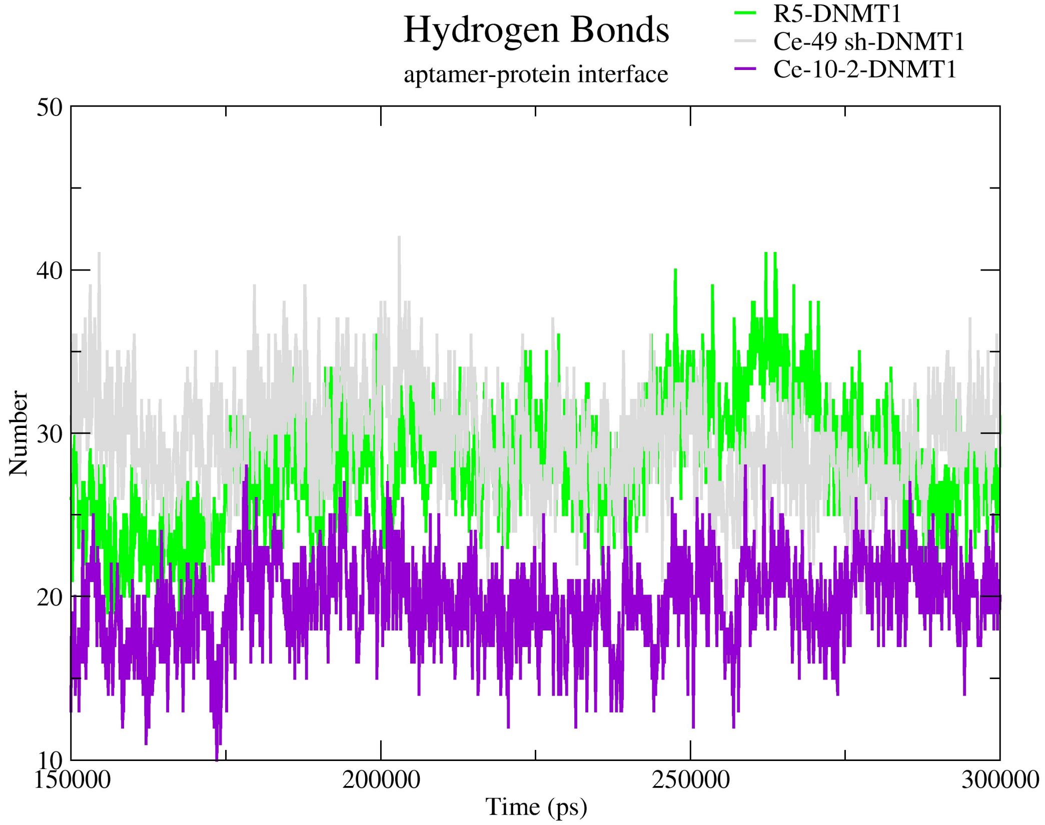


**Supplementary Figure 8. Hydrogen at complexes interfaces during the last 150 ns.** Absolute number of R5- or aptamers-DNMT1 interfaces hydrogen bonds during the last 150 ns of simulation time. The following colour code was used: R5-DNMT1: green, Ce-49 sh-DNMT1: gray and Ce-10-2 sh-DNMT1: violet.

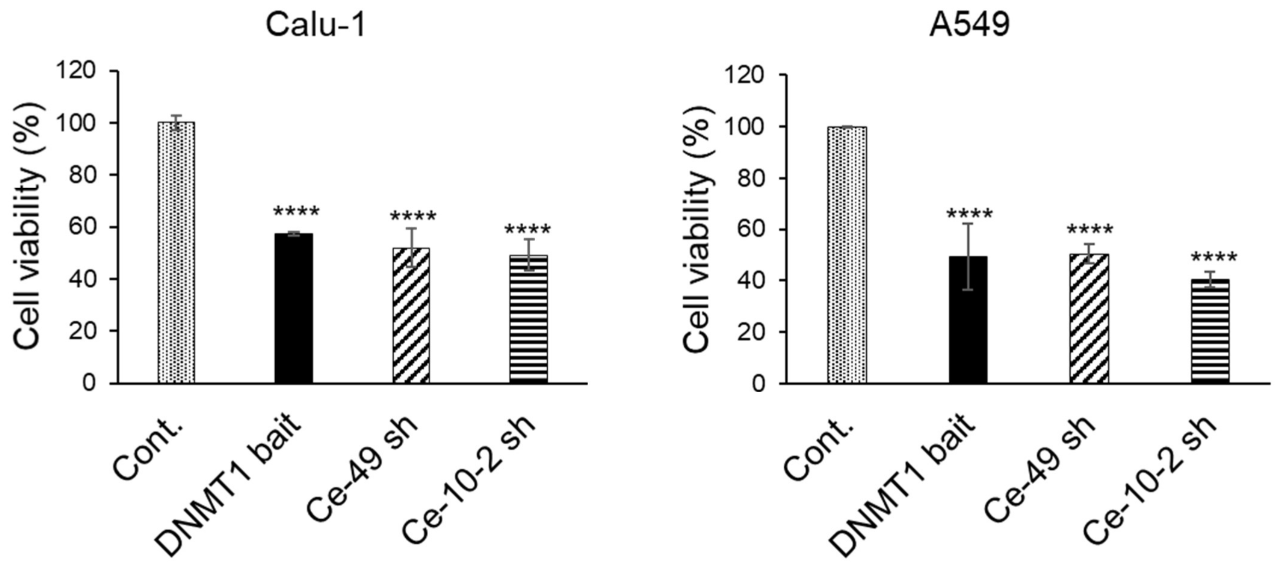


**Supplementary Figure 9. Cell viability on NSCLC cells**. Cell viability of indicated NSCLC cells transfected with indicated aptamers or Cont. for 72 hours. Error bars depict mean ± SD. Statistics by one-way Anova: ****, $p<0.0001$

**Supplementary Table 1. SELEX conditions**

| Round | RNA pool (pmol) | Protein (pmol) | RNA: protein ratio | Number of washes | Number of counter-selection |
|---|---|---|---|---|---|
| 1 | 300 | 10 | 30:1 | 2 | 1 |
| 2 | 150 | 5 | 30:1 | 3 | 1 |
| 3 | 150 | 5 | 30:1 | 4 | 1 |
